# Mitochondrial thermogenesis regulates heat-shock response in the nucleus

**DOI:** 10.1101/2023.12.18.571173

**Authors:** Hee Yong Lee, Hwa-Ryeon Kim, Chulhwan Kwak, Myeong-Gyun Kang, Jae-Seok Roe, Hyun-Woo Rhee

**Affiliations:** Department of Chemistry, Seoul National University, Seoul 08826, Korea; Department of Biochemistry, Yonsei University, Seoul 03722, Korea; School of Biological Sciences, Seoul National University, Seoul 08826, Korea; Department of Neurosurgery, Stanford University, Stanford, CA, 94305, USA

## Abstract

Mitochondrial thermogenesis is a process in which heat is generated by mitochondrial respiration. In living organisms, the thermogenic mechanisms that maintain body temperature have been studied extensively in fat cells, with little knowledge on how mitochondrial heat may act beyond energy expenditure. Here, we highlighted exothermic oxygen reduction reaction (ΔHf° = -285 kJ/mol) is the main source of the protonophore-induced mitochondrial thermogenesis and this heat was conducted to other cellular organelles, including the nuclei. As a result, mitochondrial heat that reached the nucleus initiated the classical heat shock response, including the formation of nuclear stress granules and localization of heat shock factor 1 to chromatin. Consequently, activated HSF1 increases gene expression associated with the response to thermal stress in mammalian cells. Our results illustrate heat generated within the cells as a potential source of mitochondrial-nucleus communication and expand our understanding of the biological functions of mitochondria in cell physiology.

## Introduction

Temperature induces diverse biological events, including biochemical reactions (e.g., reaction equilibrium and reaction rates)[1-2] and structural changes in proteins[3] and lipid membranes[4]. For this reason, living systems can precisely sense and respond to temperature changes[5]. Under the external heat stress conditions, heat shock factor 1 (HSF1) senses the increase in temperature and undergoes a structural change that enables it to regulate the expression of heat shock proteins (HSPs) thereby promoting cell survival. Although HSF1 activation under external heat shock conditions was discovered more than 30 years ago[6-9], whether mammalian cells can produce their own heat to initiate an HSF1-mediated heat shock response remains unknown.

Mitochondrial thermogenesis is an intracellular event that actively generates heat to maintain body temperature. In fat cell mitochondria, protons are directly imported into the mitochondrial matrix by the proton transporter protein (UCP1), not by ATP synthase[10], which is known to accelerate exothermic oxygen reduction to water reactions[11-12]. The similar event can be induced by treatment with mitochondrial uncoupling agents such as carbonyl cyanide *p*-(tri-fluromethoxy)phenyl-hydrazone (FCCP) which can directly import protons to the mitochondrial matrix[13]. Intriguingly, the FCCP-driven proton import to the mitochondrial matrix can also initiate mitochondrial thermogenesis which was validated experimentally using several fluorescent thermometers that can precisely detect mitochondrial temperature increases in live cells[14-21]. However, it remains uncertain what the biological importance and impact of the heat generated in mitochondria by changes in proton import is. In this study, we highlighted that exothermic proton-coupled oxygen consumption reaction or oxygen reduction reaction (ORR, O2 + 4H+ + 4e-→ 2H2O, ΔHf° = -285 kJ/mol)[22-23] (**Fig. 1A**) is the main thermogenic source under the protonophore (i.e. FCCP) treatment by regulation of oxygen concentration and electron flows in the electron transport chain. Furthermore, we determined that this ORR-induced thermogenesis in mitochondria leads to thermal conduction to the other organelles such as the nucleus. Under the same conditions, we observed that mitochondrial thermogenesis activated canonical nuclear heat shock response programs mediated by HSF1. Overall, our study reveals a cell-intrinsic mechanism that allows heat to actively convey biological signals from the mitochondria to the nucleus.

**Figure 1.**
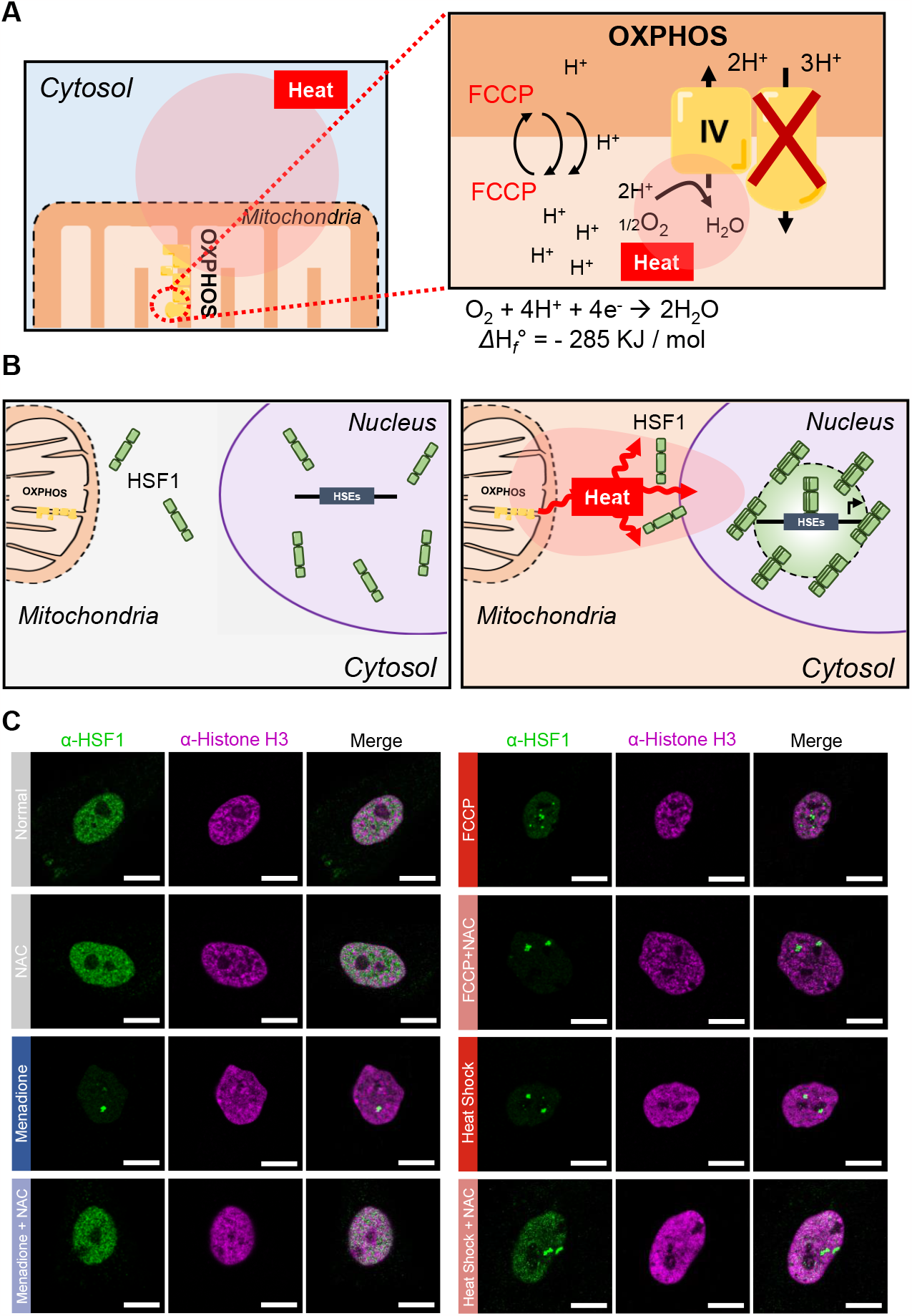
Nuclear HSF1 activation by mitochondrial thermogenesis under FCCP treatment. (**A**) Schematic representation of FCCP-induced exothermic mitochondrial oxygen reduction reaction (ORR). (**B**) Schematic representation of HSF1 activation via mitochondrial thermogenesis following FCCP treatment (HSEs: heat shock elements). (C) Confocal images of endogenous HSF1 (anti-HSF1) and Histone H3 (anti-histone H3) activation in MCF10A cells incubated with either FCCP (100 μM, 1 h) or menadione (30 μM, 30 min), or subjected to heat shock (43 °C, 1 h), and with or without co-treatment with NAC (5 mM, 1 h). Scale bar 10 μm.

## Results

### Heat shock factor 1 activation by mitochondrial thermogenesis

To determine whether the activity of HSF1 is regulated by mitochondrial heat (Fig. 1B), we checked endogenous HSF1 localization by immunofluorescence imaging following treatment with FCCP. Biochemically, the FCCP is a weak acid that enforces reversible proton import from the mitochondrial intermembrane space to the matrix, which in turn uncouples the electron transport chain and inhibits ATP synthesis[24]. From a chemical perspective, this event can induce a highly exothermic oxygen reduction reaction (ORR) to water at mitochondrial complex IV (O2 + 4H+ + 4e-→ 2H2O, ΔHf° = -285 kJ/mol)[22] by providing essential protons for this reaction (**Fig. 1A**). We applied FCCP to HEK293T cells and measured the mitochondrial inner membrane potential with TMRE (***SI Appendix*, Figs. S1A-B**). We validated that ER membrane temperature was increased by the FCCP treatment with an ER membrane-localized temperature-measuring fluorescent probe, ERthermAC (ETAC)[25](***SI Appendix*, Figs. S1C-D**). Using fluorescent polymeric thermometer (FDV) which are evenly distributed in entire cellular area[26-28], we also observed that fluorescence emission intensity ratio (FI580/FI515) of FDV was changed to the ratio at the 39 °C under the FCCP treatment (***SI Appendix*, Figs. S1E-G**).

HSF1 is the fundamental heat-sensitive nuclear factor that can be activated with nuclear “foci” formation under heat stress[29-30]. These observations collectively indicate that the mitochondria-generated heat induced by FCCP can be transferred to the surrounding space and proximal organelles. Under this condition, endogenous HSF1 formed foci in the nucleus after FCCP treatment, and this pattern was comparable to that of HSF1 foci formation under external heat shock conditions (**Fig. 1C**).

Including heat, there are several factors that contribute to the process of HSF1 activation, including intracellular pH changes[31-33] and ROS generation[34-35]. Hence, we decided to investigate whether mitochondrial heat is the primary factor that triggers HSF1 foci formation with FCCP treatment. First of all, we excluded pH changes in mediating HSF1 activation as FCCP treatment showed no changes in nuclear pH measured by pHluorin2[36] (***SI Appendix*, Fig. S2A**). To assess ROS generation under the FCCP treatment, we employed our recently developed system based on the ROS-dependent engineered ascorbate peroxidase (APEX) reaction[37] (**Fig. 2**). Notably, the ROS production induced by FCCP was significantly lower in all submitochondrial spaces compared to the well-established ROS-generating agent, menadione[38] (**Figs. 2D-E**).

**Figure 2.**
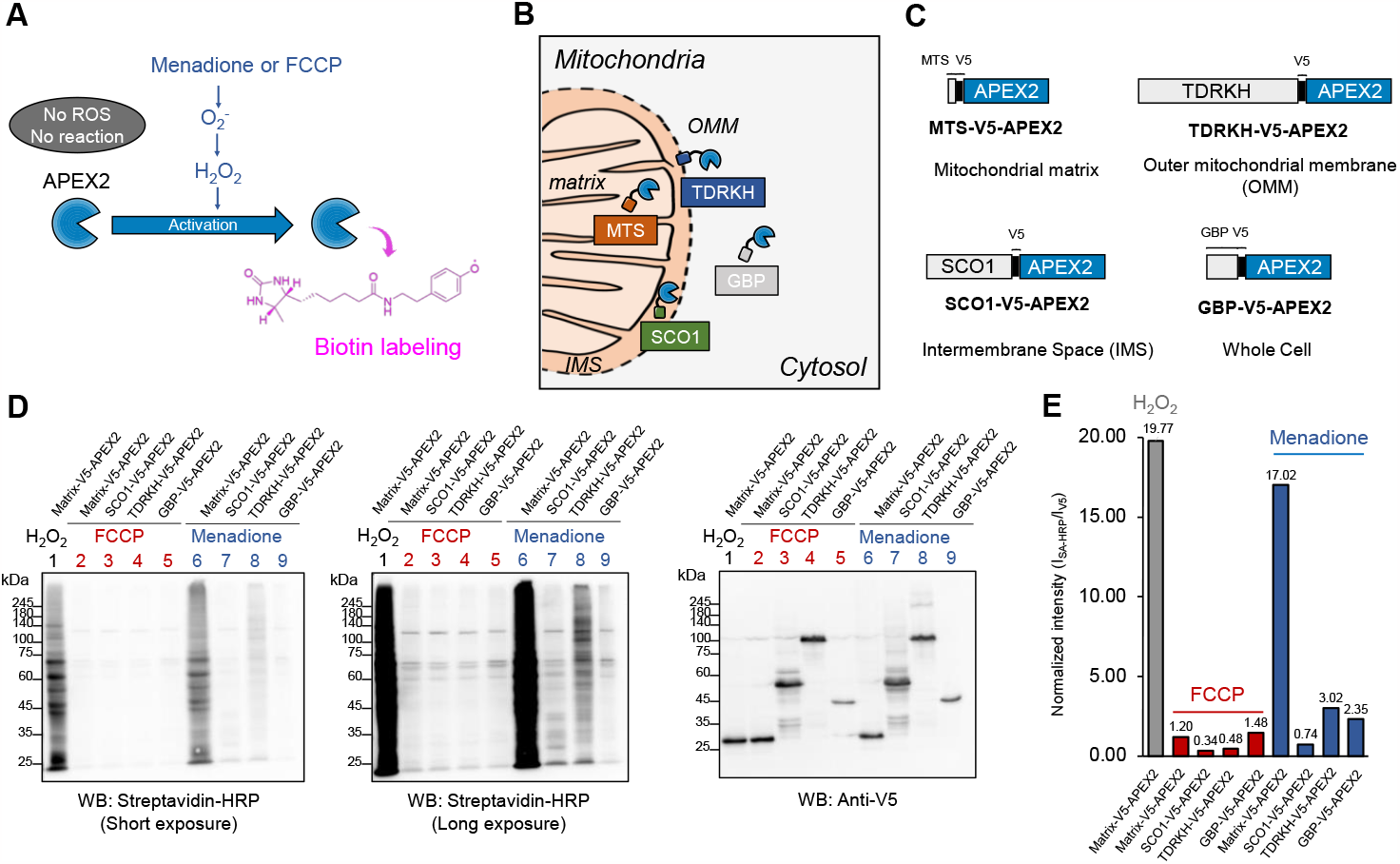
FCCP treatment does not induce ROS generation. (**A**) Schematic depiction of APEX-mediated recording of cellular ROS generation. (**B**) Graphical representation of the subcellular localization of APEX2 constructs used for the enzymatic recording of hydrogen peroxide generation: Matrix-V5-APEX2 (mitochondrial matrix), SCO1-V5-APEX2 (IMS), TDRKH-V5-APEX2 (OMM), and GBP-V5-APEX2 (whole cell). (**C**) Construct map of the APEX plasmids used in this study. (**D**) Streptavidin (SA) western blot results of APEX-mediated biotinylating activity measurements after menadione or FCCP treatment. Anti-V5 western blotting images of the same lysate are shown below. (**E**) Quantification plots of the relative band intensity after short exposure to streptavidin and anti-V5 antibodies.

Notably, HSF1 has multiple activation modes either by heat stress or by ROS generation[39-40]. We confirmed that HSF1 foci formation was induced by menadione without heat induction, however, this ROS-induced HSF1 activation by menadione was significantly quenched by co-treatment of the ROS quenching agent N-acetylcysteine (NAC, 5mM) (**Fig. 1C**). In contrast FCCP-driven HSF1 foci formation was minimally affected by co-treatment with the same concentration of NAC (5mM), suggesting that ROS is unlikely attributed to HSF1 foci formation (**Fig. 1C**). These results collectively demonstrate that FCCP-induced HSF1 foci formation is independent of ROS or pH changes, and instead, it can be attributed to mitochondrial thermogenesis.

Motivated by the above results, we investigated the potential use of HSF1 foci formation as a real-time recording system that responds to heat conducted from mitochondria. To facilitate the observation of HSF1 foci formation, we prepared an HSF1-EGFP construct that can be delivered lentiviral and expressed stably in cells (**Fig. 3A**). Utilizing the GFP fluorescence, we recorded the HSF1-EGFP foci formation immediately following FCCP treatment by real-time confocal microscope (Fig. 3B; movie 1-2). In this real-time experiment, cells were incubated in normal growth media without FCCP treatment for the first 10 min and we did not observe any HSF1-EGFP foci in cells. However, when exposed to FCCP, HSF1-EGFP foci emerged immediately and disappeared rapidly when replaced with FCCP-free growth medium (**Figs. 3B-C**). The time-course measurement of HSF1-EGFP foci number exhibited a strong positive correlation with the presence or absence of FCCP in the cell growth medium (**Fig. 3C**). This result also presented that the HSF1 foci can be induced in a reversible manner by transient heat generation. Consistently, HSF1-EGFP foci formation was also successfully promoted by other mitochondrial uncouplers such as BAM15 and CCCP (Carbonyl cyanide 3-chlorophenylhydrazone)[24] (***SI Appendix*, Figs. S2B-E**). Nuclear HSF1 foci formation following FCCP treatment was again detected to a comparable degree in multiple other cell lines, including U2OS, HEK293T, MCF10A, and A549 (***SI Appendix*, Figs. S2F-I; movie 3-10**). These observations indicated that there is no cell-type specification for the induction of HSF1 foci by FCCP. Taken together, our collective results suggest that HSF1 activation via mitochondrial thermogenesis is conserved in mammalian cells.

**Figure 3.**
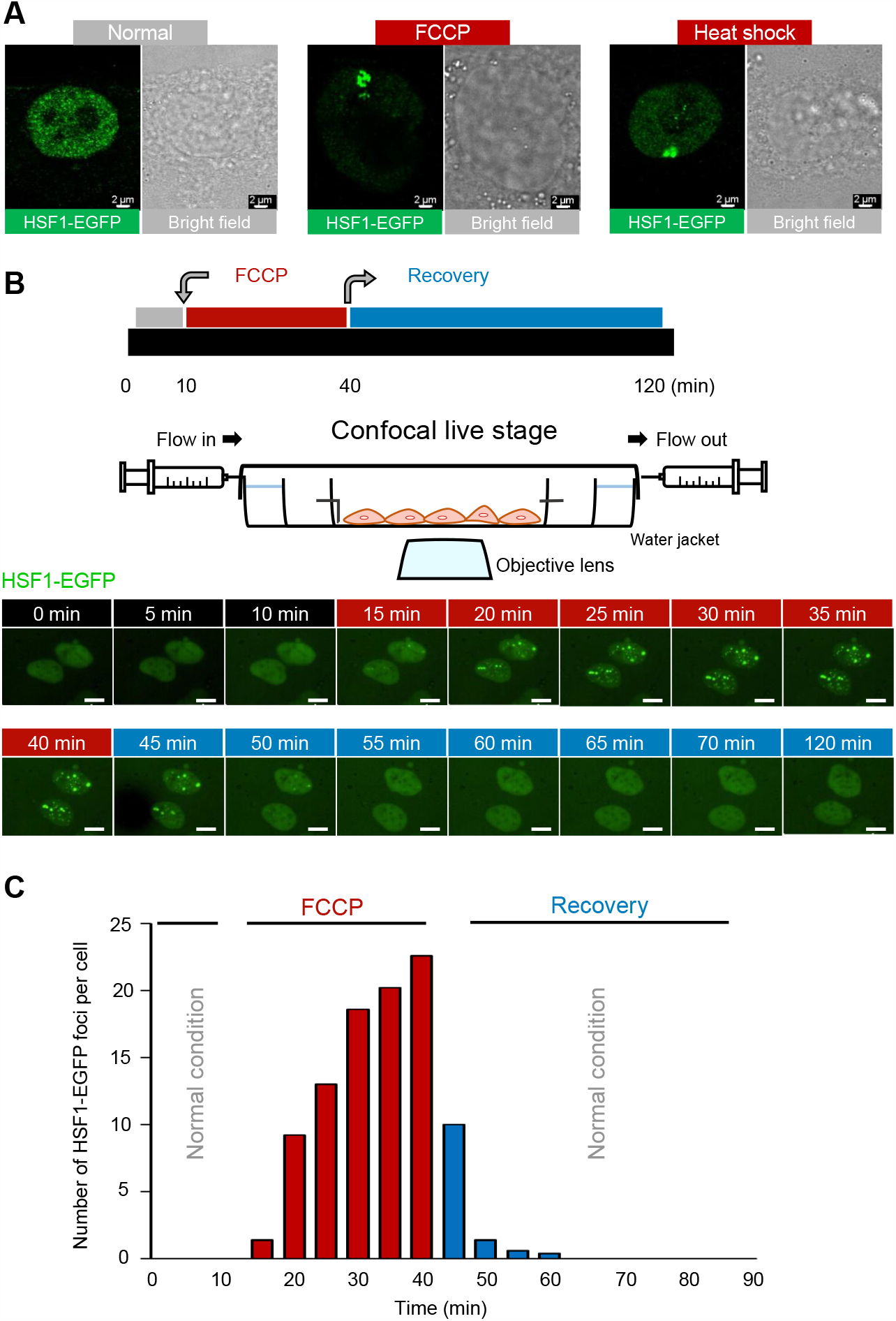
Real-time imaging of HSF1 foci formation by FCCP treatment. (**A**) Confocal fluorescent images of HSF1-EGFP foci formation in the stable HSF1-EGFP-expressing cells, under FCCP treatment or heat shock. Scale bar 2 μm. (**B**) Real-time fluorescence recording of HSF1-EGFP in U2OS HSF1-EGFP-expressing stable cells. The FCCP (100 μM, 30 min) was administered after 10 min of incubation in normal conditions, and recovery was recorded during the incubation in fresh media for 1 h. (C) Time-series graph for the number of counted HSF1-EGFP foci per cell nucleus throughout the FCCP-treatment and recovery phases.

### Exothermic oxygen consumption reaction drives the HSF1 foci formation

Mitochondrial thermogenesis, induced by exothermic oxygen reduction reaction (ORR, O2 + 4H+ + 4e-→ 2H2O, ΔHf° = -285 kJ/mol)[22], relies on the abundant availability of reactants (H+, O2, and electrons) within the mitochondria. Consequently, protonophore-induced mitochondrial thermogenesis is contingent upon well-maintained oxygen supply and efficient electron transport in the mitochondria. Thus, we hypothesized that either diminished oxygen levels or impaired electron transport during FCCP treatment may weaken heat generation thereby attenuating formation of HSF1 foci (**Fig. 4A**).

**Figure 4.**
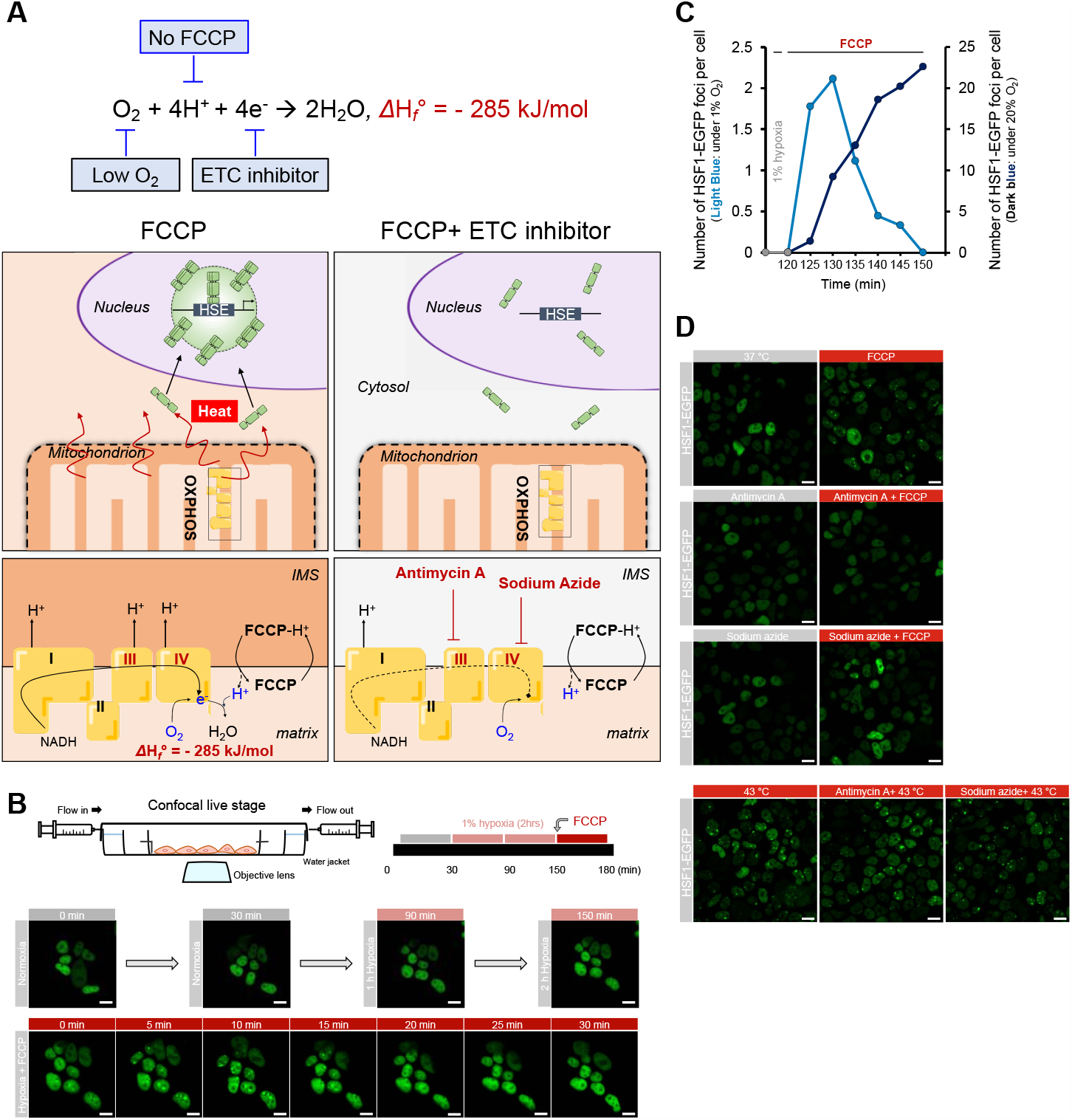
Mitochondrial oxygen concentration and OXPHOS complex activity are crucial for mitochondrial thermogenesis and HSF1 activation. (**A**) Schematic view of exothermic mitochondrial oxygen consumption reaction under the FCCP treatment. (**B**) Real-time confocal imaging of HSF1-EGFP stably expressed in HEK293T cells under normoxic (20% O2 concentration, 30 min) and hypoxic (1% O2 concentration, 2 h 30 min) conditions, with FCCP (100 μM) co-treatment at last 30 min. Scale bar 10 μm. (**C**) The number of counted HSF1-EGFP foci per cell under incubation conditions of 100 μM FCCP and either 1% O2 concentration (Light blue dots, count: left y-axis), or 20% O2 concentration (Dark blue dots, count: right y-axis). The gray dots represent 1% hypoxia without any chemical treatment. (**D**) Confocal images of HSF1-EGFP activation in HEK293T HSF1-EGFP stably expressed cells treated with FCCP (100 μM, 1 h), antimycin A (20 μM, 1 h), and sodium azide (32 μM, 1 h). Scale bar 10 μm. (**E**) Confocal images of HSF1-EGFP activation in HEK293T HSF1-EGFP stably expressed cells treated with antimycin A (20 μM, 1 h), sodium azide (32 μM, 1 h), under heat shock (43 °C, 1 h) conditions. Scale bar 10 μm.

To test this hypothesis, we reduced oxygen supply decreasing the availability of oxygen molecules in the media and the intracellular environment (**Fig. 4B; movie 11**). The HSF1-EGFP expressing cells were subjected to 2 h of hypoxic conditions (1% O2) prior to incubation with 100 μM FCCP. Using a real-time confocal imaging microscope, we could not detect HSF1 foci formation under prolonged hypoxia of over 2 h (**Fig. 4C**; gray dots), indicating that 2–3 h of hypoxia are sufficient to attenuate FCCP-dependent HSF1 foci formation. We observed only 1–2 HSF1 foci per cell within 10 min of FCCP treatment; however, these foci subsequently disappeared in the span of 30 min (**Fig. 4C**; light blue dots). In contrast, under normoxic baseline conditions (20% O2), approximately 10 foci per cell formed within 10 min following FCCP treatment (**Fig. 4C**; dark blue dots). This finding highlights that oxygen supplementation is a pre-requisite for both mitochondrial thermogenesis and HSF1 activation.

Next, to determine if HSF1 foci formation is affected by preventing electron supply to the exothermic ORR reaction (O2 + 4H+ + 4e-→ 2H2O, ΔHf° = -285 kJ/mol), HSF1-EGFP stably expressed cells were treated with antimycin A or sodium azide, which inhibit electron flow in the Oxidative Phosphorylation (OXPHOS) complex III and IV, respectively (**Fig. 4A**). As expected, we observed that HSF1 foci formation was significantly reduced upon co-incubation of cells with OXPHOS inhibitors and FCCP (**Fig. 4D**). Conversely, HSF1-EGFP foci were distinctly formed in the control group after external heat shock at 43 °C, regardless of treatment with either of the OXPHOS inhibitors (**Fig. 4E**). This result confirmed that OXPHOS inhibitors specifically blocked the mitochondrial thermal activation process, not touching the HSF1 itself. Overall, here we confirmed that every component (i.e., proton, oxygen, electrons) in the exothermic mitochondrial ORR required for generating sufficient mitochondrial heat that can activate HSF1.

#### Mitochondrial thermogenesis activates HSF1-dependent transcriptional programs

Since heat-induced phosphorylations are prerequisites for the translocation and activation of HSF1 in the nucleus[41-43], we evaluated whether mitochondrial thermogenesis induced by FCCP treatment induces phosphorylation on HSF1. As shown in the HSF1 western blot results (***SI Appendix*, Figs. S3A-D**), HSF1 molecules moved more slowly in the gel when treated with FCCP, similar to when the heat shock is generated externally[41]. In contrast, relative to the control group, HSF1 showed no different in-gel migration under intracellular ROS-generating conditions induced by either menadione or rotenone treatment (***SI Appendix*, Figs. S3A-D**). These results implicate that phosphorylation may occur on HSF1 during FCCP-dependent mitochondrial thermogenesis. Taken together, our results suggest that mitochondrial thermogenesis by FCCP may facilitate transcriptional activity of HSF1 in the nucleus through a similar mechanism by the heat applied externally.

To validate this hypothesis, we investigated whether nuclear-localized HSF1 accumulation drives the expression of heat shock response-related programs. To determine whether HSF1 binds to heat shock response genes, we mapped the genome-wide binding of HSF1 using chromatin immunoprecipitation sequencing (ChIP-seq). Meta-analysis of the ChIP-seq datasets revealed 621 sites that commonly gained HSF1 signals in the promoter regions, termed Common-GAIN regions (**Fig. 5A**). Moreover, we found sites that recruited HSF1 under either heat shock or FCCP-treatment conditions, termed HS-GAIN and FCCP-GAIN regions, respectively (***SI Appendix*, Fig. S4**). Notably, the average intensity of FCCP-mediated HSF1 binding to Common-GAIN sites was reduced to ∼50% of that compared to heat-shock dependent HSF1 activation, corroborating that FCCP treatment increased nuclear temperature to levels lower than 43 °C (**Figs. 5B-C**). This result is in good agreement with our previously measured mean temperature of entire cellular area (39 °C) using FDV under the FCCP treatment (***SI Appendix*, Fig. S1G**). The motif analysis of the Common-GAIN regions returned classical HSF1 binding motifs, although the possibility of engagement of other transcription factors could not be excluded (**Fig. 5D**). With luciferase assay system, we confirmed that HSPD1/HSPE1 promoter which is one of Common-GAIN regions (**Fig. 5C**) can be activated both external heat shock and FCCP treatment conditions (***SI Appendix*, Figs. S5A-B**).

**Figure 5.**
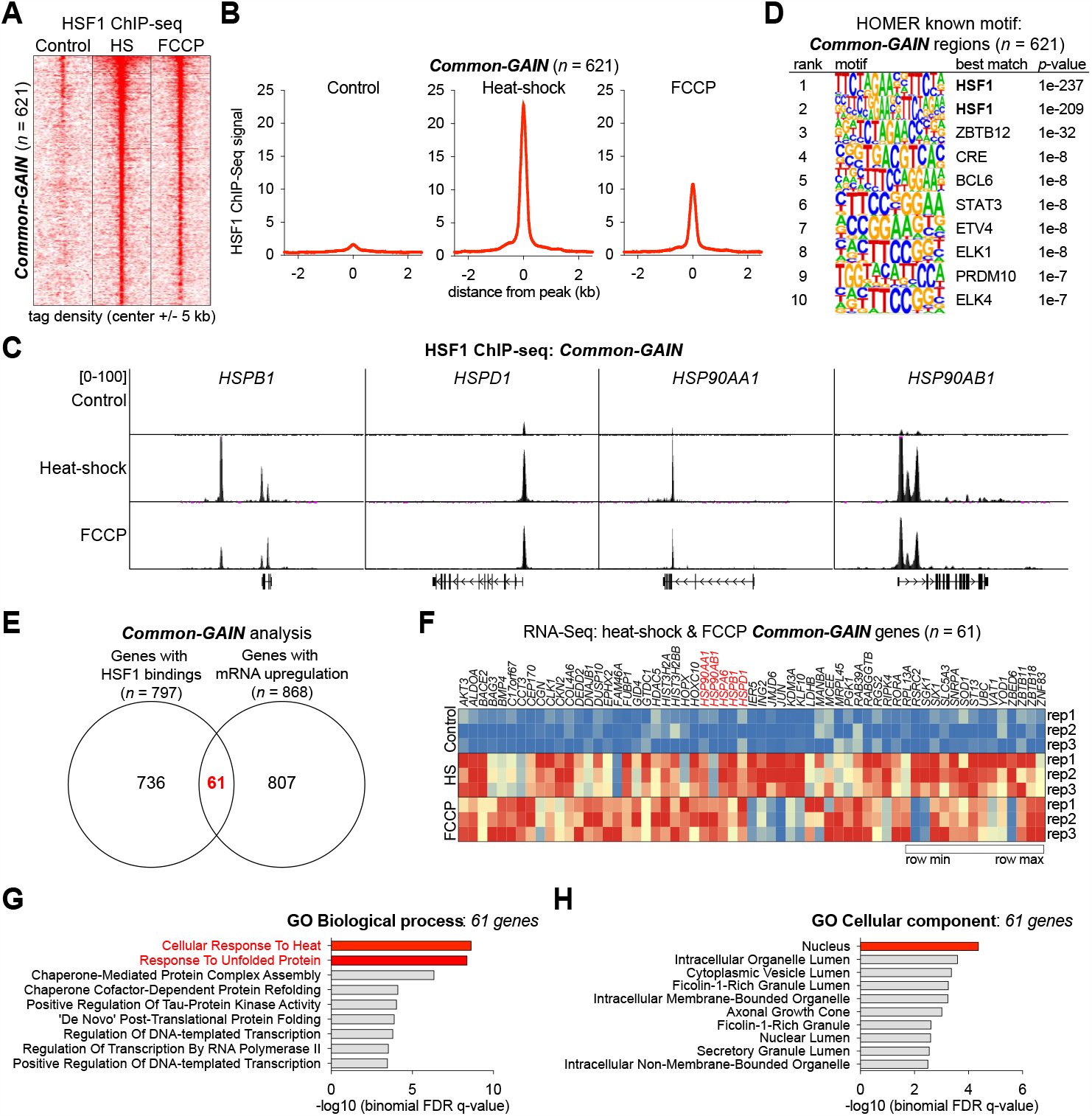
Chip-seq and RNA-seq analysis for common-GAIN regions of HSF1 expression under heat shock and FCCP treatment. (**A**) Density plots of HSF1 ChIP-seq signal enrichment at a 5-kb region around the center of heat shock/FCCP co-bound regions. Each row represents a single region (n = 621). The conditions for heat shock were 43 °C for 1 h, and for the FCCP treatment 100 μM, for 1 h. (**B**) Analysis of ChIP-seq signals on overactive promoters and enhancers under control, heat shock, and FCCP-treatment conditions. The scale of all graphs is calculated based on the distance of the peak (kb). (**C**) Representative HSF1 ChIP-seq profiles at the HSPB1, HSPD1, HSP90AA1, and HSP90AB1 loci. (**D**) Motif analysis (HOMER) based on HSF1 ChIP-seq results showing enriched motifs in Common-GAIN peaks. Only the top 10 motifs ranked by statistical significance are shown in the logo plot. (**E**) Venn diagram depicting the number of identified HSF1-bound genes (n = 797) and mRNA-upregulated genes (n = 868), following heat shock or treatment with FCCP. Between the two clusters, there were a total of 61 (n = 61) overlapping genes, identified as both being associated with HSF1 and upregulated. (**F**) Heatmap representation of changes in gene expression of heat shock and FCCP Common-GAIN genes (n = 61). (G–H) Gene Ontology (GO) analysis for the identified 61 heat shock/FCCP co-bound region genes of (G) the associated biological processes, and (H) the subcellular compartment (Cellular Component).

Among the 797 genes in which single or multiple HSF1-binding peaks were found in the promoter regions, RNA-seq analysis further revealed that 61 genes were transcriptionally activated in FCCP-treated cells to a similar extent as under external heat shock conditions (**Figs. 5E-F**). Furthermore, gene ontology (GO) analysis revealed that 61 genes were significantly associated with the nuclear response to heat shock stress, exemplified by “cellular response to heat” and “response to unfolded protein,” indicating that the overall transcriptional output of the FCCP treatment phenocopies that of the external heat shock stimulus (**Figs. 5G-H** and ***SI Appendix*, Figs. S5C-D**). Taken together, our findings suggest that FCCP-mediated heat generation resembles the biological activity of the heat shock response at the transcriptional level.

## Discussion

In this work, we used HSF1 as a molecular sensor to measure how mitochondrial heat affects binding events of proximal proteins. As a result, we found that HSF1 formed foci in the nucleus when we added protonophore to potentiate the mitochondrial reaction for heat-releasing oxygen consumption. At the same time, we also observed HSF1 foci formation, which could be reversed by controlling the oxygen consumption in the mitochondria. Finally, integrative HSF1 ChIP-seq and RNA-seq experiments verified the activation of functional HSF1 pathway in a similar way under external heat stress conditions.

Our results revealed that mitochondrial thermogenesis can modulate the activity of intracellular thermosensing proteins such as HSF1, in live cells. This suggests that mitochondria-generated heat could act as an intracellular signal that may lead to protein conformation alterations without intermediate molecular interactions. Compared to the targeted signaling by molecular transfer events (i.e., molecular conversion and translocation[44-46], molecular interactions, or modifications), mitochondrial heat can be a global retrograde signal that spatiotemporally affects numerous proteins. In mammalian cells, many proteins have a low melting temperature around 40 °C[47] and proteins with intrinsically disordered proteins (IDRs) can occur liquid-liquid phase separation (LLPS) in the range of 40 °C[48-52]. Notably, HSF1 possesses a long IDR domain (221–383 aa) that can induce LLPS[53]. Therefore, it is expected that many of such low-melting-temperature or IDR-containing proteins can be the primary effectors of mitochondrial heat signaling.

It is noteworthy that HSF1 has been studied for its role in mitochondria and nucleus communication. A group of studies have shown that HSF1 can sense mitochondrial misfolding stress and mediated mitochondrial protein unfolded response (UPRmt) in the nucleus[9],[44]. Other studies have shown that HSF1 can be activated by mitochondrial ROS and protect cells under the mitochondrial stress conditions[44, 54-55]. Our study firstly showed that mitochondrial generated heat can also activate HSF1. Our data also reinforces the hypothesis that HSF1 is a key protein that acts as a master message-transducer between the mitochondria and the nucleus. It is also noteworthy that HSF1 foci formations were observed in several aggressive tumors[8, 29]. Since high mitochondrial oxygen consumption rates have been measured in several cancer cell lines[56-58], further studies would be attempting to test whether the mitochondrial thermogenesis can promote HSF1 activation under those conditions[9, 44, 54-55] because mitochondrial temperature has not been measured in those studies.

We also provided a new chemical interpretation of mitochondrial thermogenesis, based on the exothermic property of the oxygen reduction reaction (O2 + 4H+ + 4e-→ 2H2O, ΔHf° = -285 kJ/mol), which is well-recognized in other research fields[22-23, 59]. To the best of our knowledge, our study highlights the exothermic enthalpy change of this oxygen consumption reaction (ΔHf° =-285 kJ/mol) and suggests that this is likely the primary thermogenic reaction in mitochondria. We validated that every component (i.e., protons, oxygen, and electrons) in this reaction is crucial for sufficient mitochondrial thermogenesis and the subsequent upregulation of HSF1. As the oxygen consumption rate is regarded as a reliable indirect measurement standard for thermogenic events in the metabolism research field[60], we believe that our theorem may be accepted in the field.

In summary, our current study demonstrated that mitochondrial-generated heat is sufficient to activate HSF1 independently of ROS and provides supporting evidence to consider intracellularly generated heat as a distinct signal that promotes subsequent changes in cell homeostasis. Our work may serve as a basis for future investigations to delineate the complex relationships between various mitochondrial thermogenic events and the activation of heat-sensitive proteins under diverse physiological and pathological contexts.

## Author information

### Authors

Hee Yong Lee –

Hwa-Ryeon Kim –

Chulhwan Kwak –

Myeong-Gyun Kang –

### Funding

This work was supported by funding from the National Research Foundation of Korea (NRF 2022R1A2B5B03001658, NRF-2023K2A9A2A08000203, and RS-2023-00265581 awarded to H.W.R. 2021R1A2C4001420 to J-. S.R, NRF-2022R1C1C2006983 to C. K., and NRF-2020R1I1A1A01069160 to H.R.K.). H.R.K. and J.S.R. were also supported by the Brain Korea 21 FOUR Program awarded to Yonsei University. Research conducted at the Organelle Network Research Center (NRF-2017R1A5A1015366) and the Korea Health Industry Development Institute (KHIDI) was funded by the Ministry of Health & Welfare and the Ministry of Science and Information & Communication Technology (ICT), Republic of Korea (grant number: HU23C0204). H.W.R and J.S.R are supported by Samsung Science & Technology Foundation (SSTF-BA2201-08 to H.W.R, SSTF-BA2002-12 to J.S.R).

## Notes

The authors declare no competing of interests.

## Acknowledgements

The ERthermoAc (ETAC) was a gift from Prof. Young-Tae Chang (POSTECH, Korea).

## Supporting Figure and Figure Legends

**Figure S1. Confirm the intracellular thermogenesis by two different thermometers**. (A) Chemical structures of TMRE, FCCP, and menadione. (C) Flow cytometry analyses and its histogram of HEK293T cell suspensions treated with either TMRE, FCCP, or menadione. (D) Schematic illustration of temperature measurement by ETAC. (E) ETAC fluorescence intensity after FCCP (100 μM, 1 h) treatment. Decreased ETAC fluorescence intensity under the indicates increased local temperature at the ER membrane. Flow cytometry analysis result of ETAC fluorescent signal under the two conditions (steady-state or 100 μM FCCP, 1 h), in HEK293T, HeLa and HepG2 cell lines. Cell count is shown on the y-axis, while fluorescent signal intensity (PE-A) is shown on the x-axis. (F) The polymer structure of the organic thermometer “FDV-0005”. (G) Graph of the calibration curve and the FCCP treatment results for the organic thermometer. The thermometer was used at a dilution of 0.1% w/v in cell extract solution. Measurements were performed at a temperature range of 30 °C to 45 °C. ( H) Intracellular temperature calibration by the FDV-0005 organic thermometer. The thermometer was used at a dilution of 0.01% w/v in 5% glucose solution. Measurements were performed at a temperature range of 35 °C to 45 °C. Cells were incubated at 37 °C and 5% CO 2 in a humidified incubator and treated with 100 μM FCCP.

**Figure S2. Measurement of pH under FCCP condition and HSF1-foci formation in various HSF1-EGFP stably expressed cell lines**. (A) Measurement of nuclear pH in the FCCP condition. The fluorescent protein-based pH sensor pHluorin2 was used. The HEK293T cells were incubated at 37 °C and 5% CO2 in a humidified incubator and 100 μM FCCP was used for this assay. (B-C) Chemical structures of BAM15 and CCCP. (D) Confocal images of HSF1-EGFP foci formation in HEK293T HSF1-EGFP-expressing cells following treatment with various CCCP concentrations (10, 50, and 100 μM; 1-h incubation). Scale bar 20 μm. (E) Confocal images of HSF1-EGFP foci formation in HEK293T HSF1-EGFP-expressing cells under treatment with various BAM15 concentrations (10, 20, 50, and 100 μM; 1-h incubation). Scale bar 20 μm. (F) Live-cell imaging of HSF1-EGFP foci formation following FCCP treatment (100 μM, 30 min) of HSF1-EGFP-expressing MCF10A cells. A real-time video recording is shown in movie zip 6. (G) Live-cell imaging of HSF1-EGFP foci formation following FCCP treatment (100 μM, 30 min) of HSF1-EGFP-expressing HEK293T cells. (H) Live-cell imaging of HSF1-EGFP foci formation following FCCP treatment (100 μM, 5 min) of HSF1-EGFP-expressing U2OS cells. A real-time video recording is shown in movie zip 5. (I) Live-cell imaging of HSF1-EGFP foci formation following FCCP treatment (100 μM, 30 min) of HSF1-EGFP-expressing A549 cells. All live cell experiments were conducted at 37 °C in a humidified 5% CO2 incubator.

**Figure S3. Western blot analysis in MCF10A cells**. (A-B) Western blot analysis of (A) endogenous HSF1, or (B) recombinant HSF1-EGFP modification, following treatment with FCCP (100 μM, 1 h) or heat shock (43 °C, 1 h) in MCF10A cells. Histone H3 was used as reference. (C-D) Ponceau S image of western blot results (A) and (B) respectively.

**Figure S4. Chip-seq and RNA-seq analysis of HSF1 under heat shock-specific and FCCP treatment-specific GAIN**. (A–B) Representative HSF1 ChIP-seq profiles at the (A) BAG3 and NXT2, and (B) RBM25 and AQP12B loci. (C) Density plots of HSF1 ChIP-seq, signal enrichment at a 5-kb region around the center of heat shock-specific bound regions. Each row represents a single region (n = 548). (D) Analysis of the ChIP-seq signals (shown in C) overactive promoters and enhancers. The scale of all graphs was calculated using the distance of the peaks (kb). (E) Motif analysis (HOMER) based on HSF1 ChIP-seq results showing enriched motifs in HS-GAIN peaks. Only the top 10 statistically significant motifs are shown in the logo plots. (F) Density plots of HSF1 ChIP-seq and signal enrichment at a 5-kb region around the center of the FCCP treatment-specific bound regions. (G) Analysis of the ChIP-seq signals (shown in f) overactive promoters and enhancers. The scales of all graphs were calculated using the distance of the peaks (kb). (H) Motif analysis (HOMER) based on HSF1 ChIP-seq results showing enriched motifs in FCCP-GAIN peaks. Only the top 10 statistically significant motifs are shown in the logo plots.

**Figure S5. Luciferase assay on HSPD1; HSF1 binding site and mRNA upregulated gene under Heat shock and FCCP treatment common-GAIN**. (A) HSPD1/HSPE1 promoter region sequence, and schematic representation of the HSPD1-luciferase assay. (B) HSPD1/HSPE1 promoter luciferase assay in HEK293T cells after FCCP treatment (100 μM, 1 h), heat shock at 40 °C for 1 h, or heat shock a 43 °C for 1 h). All live cell experiments were conducted at 37 °C, in a humidified 5% CO2 incubator. Box plots shown in b indicate the quartiles, whiskers range from minimal to maximal values except outlier. The middle line of the box indicates the mid-point of the data (middle quartile). Each dot indicates individual data points plotted on the middle side of the box. Symbol ‘x’ means average value of data. Statistical analysis performed by student’s T test and the following symbols were used:(**p<0.01). (C) Venn diagram showing the overlap (n = 15) of the HSF1-bound genes (n = 708) and mRNA upregulated genes (n = 104) under heat shock-specific conditions. (D) Gene Ontology (GO) analysis of the biological processes associated with the identified 15 upregulated HSF1-bound genes (as shown in c).

## Notes

### Competing Interest Statement

The authors have declared no competing interest.

