## Supporting information and supplemental figures 1 to 5 for "Mitochondrial thermogenesis regulates heat-shock response in the nucleus"

### Table of contents:

Materials and methods ............ S2-9

Plasmid information ............ S10

Figure S1 ............ S11

Figure S2 ............ S12

Figure S3 ............ S13

Figure S4 ............ S14

Figure S5 ............ S15

Supplemental Movie legends............ S16

Reference ............ S17

**Materials and Methods**

**Plasmids and Cloning**. Genes were cloned into appropriate vectors using standard enzymatic restriction digestion and ligated with the T4 DNA ligase. The PCR products were digested with the same restriction enzymes and ligated into cut vectors (e.g., pcDNA5, and pFUGW). For pcDNA5, the CMV promoter was used, whereas for pFUGW, the UBC promoter was used for expression in mammalian cells. Plasmid information depicts the gene constructs used in this study.

**Cell Culture**. The HEK293T cells were obtained from ATCC (Manassas, VA, USA), while the U2OS and A549 cells were obtained from the Korean Cell Line Bank. All cell lines were cultured according to standard mammalian tissue culture protocols at 37 °C and 5% $CO_2$ in a humidified incubator. Cell lines at 60–80% confluence was transiently transfected with polyethyleneimine (PEI, Polysciences, 23966). The MCF10A cell line (ATCC-CRL-10317) was a gift from Mi-Ock Lee (Seoul National University) and cultured in DMEM / F12 (1:1) (1X) (Gibco, 11330-032) supplemented with 5% FBS, 20 μg/ml EGF (Peprotech, 1 mg), 0.5 mg/ml hydrocortisone (Sigma, H-0888-1g), and 10 μg/ml insulin (I9278-5ML).

**Transfection and Construction of Stable Cell Lines**. In this study, we generated stable cell lines via lentiviral infection or Flp-In system. For the generation of lentivirus, the HEK293T cells were cultured in DMEM (Corning) supplemented with 10% FBS, at 37 °C and 5% $CO_2$. The cells were grown in a 6-well cell culture plate containing 2 ml of media. Lentivirus were generated by co-transfection of 60–80% confluent cell cultures using PEI (Polysciences, 23966) and vector plasmids. The transfection per well was performed with approximately 1000 ng plasmid DNA mixed with 4 μg of PEI (Polysciences, 23966), 375 ng of pMDL.g/pRRE (Addgene # 12251), 375 ng of pRSV-REV(Addgene # 12253) or 250 ng of pMD2.g (a gift from Jiyoung Park in UNIST), and each diluted in 200 μL DMEM without FBS. Following a 2–3 h incubation, the medium was replaced with complete DMEM to allow the production of the recombinant lentivirus in transfected cells. All media containing the recombinant virus were collected 24 h after transfection and new complete DMEM was added to the cell culture. Each collected virus-containing medium was placed into a 15 ml tube and stored at 4 °C. Virus media were filtered through syringe-driven filters (13 mm, PES membrane, 0.45 μM, sterile, J1.F404.013N, JetBiofill) and stored at 4 °C for a week, or at -80 °C for a month. Cell lines (HEK293T, U2OS, A549, MCF10A) were cultured in a 6-well cell plate under appropriate conditions and when at 60% confluence, the medium was replaced with the filtered virus medium. Cell cultures were incubated for a further 3 h for MCF10A, and 16 h for HEK293T, U2OS, and A549 cells. Subsequently, the medium was changed and scaled up to the desired experimental volume. For stable cell line generation via Flp-In system, the Flp-In™ T-REx™ 293 cells (Life Technologies) were cultured in DMEM (Corning) supplemented with 10% FBS, at 37 °C and 5% $CO_2$. Cells were grown in T25 flasks. Stable cell lines were generated by co-transfection with the pcDNA™5/FRT/TO expression construct and the pOG44 plasmid. Cells were transfected at 60–80% confluence using PEI (Polysciences, 23966), at a dose of 4 µl PEI and 3,600 ng of plasmid (at a 9:1 ratio of pOG44:pcDNA5) per T25 flask. After 24 h, the cells were split into a 90-mm cell culture dish (SPL, 11090) with the appropriate concentration of hygromycin B (100–200 g/mL). The medium containing hygromycin B was changed every three to four days. After 2–3 weeks, we selected between three and four colonies to transfer to 24-well plates. Cells were continuously split into larger plates, and a cell stock was prepared. After splitting the cells into 6-well plates, separate samples were prepared for expression testing. To induce protein expression in stable cells, cells at 60–80% confluence were treated with 100 ng/mL doxycycline (Sigma Aldrich). The cells were dethiobiotin-phenol-labeled and lysed 18–24 h after induction.

**Confocal Imaging and Analysis**. To visualize the subcellular localization of the transiently expressed POI, cells were plated on coverslips (thickness no. 1.5 and radius: 18 mm). For fixed cell imaging, cells were fixed with 4% paraformaldehyde and permeabilized with 0.1% Triton X-100 (Sigma, Cat# 9536-19-5) in PBS for 5–10 min at RT. Next, cells were washed with Dulbecco's phosphate-buffered saline (DPBS) and blocked for 1 h with 2% BSA in DPBS, at RT. Immunolabeling was conducted in a blocking solution with primary antibodies and Alexa-labeled secondary antibodies (mouse anti-Alexa 647 and rabbit anti-Alexa 488, 1:2000 dilution), with three times washes. Immunofluorescence images were obtained and analyzed using an SP8 X Leica microscope (NICEM, Seoul National University, Seoul, Republic of Korea) with an objective lens (HC PL APO 100x/1.40 OIL), a white light laser (470–670 nm, 1 nm tunable laser), and a HyD detector, which was controlled using the LAS X software and FV3000 microscope (Oylmpus) with an objective lens (UPLXAPO 100XO), which was controlled using the Olympus Fluoview software. The primary antibodies used for Immunofluorescence imaging were anti-HSF1 (1:500 dilution, CST, Cat# 4356) and anti-Histone 3 (1:3000 dilution, Santa Cruz, sc-517576). Secondary antibodies were Alexa Fluor-conjugated anti-mouse 647(Cat# A-21235) and anti-rabbit 488 (Cat# A-11008) purchased from Invitrogen.

**Flow Cytometry Analysis for Mitochondrial Membrane Potential and Temperature Measurement by ETAC**. Cells were seeded in 6-well plates for flow cytometry analysis. The next day, cells were incubated for 30 min with 200 nM TMRE (Cat #115532-52-0, Sigma) and 0.5 µM ETAC (a gift from Young-Tae Chang. Kriszt, Rókus, et al. *Scientific reports,* 2017). Cells were then washed three times with the growth medium and incubated overnight in fresh medium. The cells stained by each dye were further incubated with either 100 µM FCCP for 1 h, 30 µM menadione for 30 min, 1.5 µM oligomycin for 24 h, or 1 µM rotenone for 24 h. A steady state indicates the absence of chemical treatment. For the following flow cytometry analysis, cell suspensions were prepared by trypsin-EDTA (0.25%)-based cell detachment and dilution with 4× fresh medium. Samples (10000 cells/events) were examined using a FACSCanto II (BD Bioscience) or flow cytometry LX (Beckman Coulter) according to their fluorescence signals. All flow cytometry data were processed using the FlowJo 10 or CytExpert software.

**Intracellular Temperature Sensing by Cellular Thermoprobe 'FDV'.** To use FDV (Funakoshi, Cat# FDV-0005), reconstitutive 200 μg powder of FDV in 20 μl of ultrapure water to prepare a 1% w/v stock solution right before each experiment with protecting from light. For the calibration curve, cell pellets (8.8 x $10^6$) were collected from 100 mm cell culture dish and resuspended in hypertonic buffer (2.5 ml, containing 0.42 M KCl, 50 mM HEPES-KOH, 5 mM MgCl2, 0.1 mM EDTA, 20% glycerol, pH 7.8). Lyse the cells using a syringe and centrifuged at 15,000 × *g* for 15 min at 4 °C, and the supernatants were collected. Dilute the supernatant with ultrapure water up to 40% to adjust its KCl concentration to 0.15 M. Dilute 1 μl of 1% w/v FDV stock solution with cell extract (100 μl). Put the solution on a 96-well black plates (SPL, Cat. No. 30296). The fluorescence was measured using a Synergy ™ H1 microplate reader (BioTek Instruments, Inc.). A wavelength of 458 or 473 nm was used for excitation. The emission range were 490-530 nm and 570-610 nm. Measurements were taken from 3 samples, and the average value was used for the analysis. The temperature was raised by 5 °C, with a 10 min wait time to allow for solution stabilization before obtaining the next measurement. The experiment ended at a final temperature of 45 °C. All experiments began with an appropriate temperature set 10 min prior to the experiment. For the live cell temperature measurements, Cells were cultured on 96-well black plates (SPL, Cat. No. 30296) using the same conditions as described in "Cell Culture". The media volume was 200 μl for each well. To measure the fluorescence, we used cell cultures at 70–80% confluence and the initial microplate reader temperature was set at 25 °C. Although the measurement of fluorescence was same as calibration method, for the FCCP treatments, cells were used at 70–80% confluence, and the initial medium was replaced with an equal volume of medium containing the appropriate compound. Cells were incubated for 10 min, upon which time, we measured the fluorescence on a microplate reader set at a temperature of 37 °C. For data analysis, the ratio of FDV and the temperature change were estimated from the formula of the fitted standard curve that was created based on the measurements per 5 °C increase, as in that range the fluorescence intensity ratio experienced an almost linear rate of increase. Using this curve, we input the chemical condition fluorescence intensity ratio and estimated the temperature variation before and after treatment with chemical compound.

**pHluorin2 Fluorescence Measurement.** Excitation scans were performed a Synergy ™ H1 microplate reader (BioTek Instruments, Inc.). Standard HEK293 cells were plated on 96-well black plates (SPL, Cat. No. 30296) and transiently transfected with 1000 ng pHluorin2 plasmid DNA mixed with 4 μg of PEI (Polysciences, 23966). After 24 hours transfection, cells were washed with Dulbecco's phosphate-buffered saline (DPBS). Plates were placed into the microplate reader for a 10 min incubation at 37°C and measured. A wavelength of

520 nm was used for excitation. The emission range were 300-490 nm with a 2 nm step. Measurements were taken from 3 samples, and the average value was used for the analysis. For the pH standard curve, calibration buffer contained the following: 50 mM MES buffer (pH 5.5, 6.0, and 6.5), 50 mM Tris·HCl (pH 7.0, 7.5, 8.0 to 8.5).

**Live Cell Imaging and Analysis**. To image live cells, the cells were cultured in a 30 mm glass bottom confocal dish (SPL, Cat# 100350) or 18mm glass (MARIENFELD, HSU-0111580) with poly-L-lysine solution (Sigma, P8920-100ML) coating . Using an ExFluorer live cell imager (Live Cell Instrument, Namyangju, Republic of Korea). For HSF1-EGFP, the FITC channel (488 nm) was used. The confocal dishes containing the cultured cells were fastened tightly without a dish cover and placed on a stage-on incubator, maintained at 37 °C and 5% $CO_2$ in a humidified environment. For imaging the HSF1-EGFP, exposure time was set at 200 ms. The initial volume of the medium was 2 ml. For the FCCP treatment, 1 ml of medium was carefully removed from the cell culture dish and replaced with 1 ml of 2xFCCP-containing medium. For cell recovery, we replaced 1.5 ml of medium from the cell culture dish with an equal volume of fresh culture medium, before subsequently replacing the entire total volume of medium in the cell culture with new fresh culture medium. The gTEMP fluorescence intensity values were obtained using an ExFluorer analyzer (Live Cell Instrument, Namyangju, Republic of Korea), and data were exported from the imager for further analysis. The treatment and analysis of FCCP and the cell recovery method were the same as in with the gTEMP imaging experiments.

**Live Cell Imaging for FCCP Treatment-Recovery with or without Hypoxia.** To image real-time confocal imaging of HSF1-EGFP in various cell lines, cells were cultured same methods with 'Live Cell Imaging and Analysis'. A 'Incubator T Series' (Live Cell Instrument, Namyangju, Republic of Korea) is on the FV3000 microscope (Oylmpus) stage, which precisely control the gas, temperature, and humidity of the stage incubator. For imaging, all cell lines were cultured according to standard mammalian culture protocols at 37 °C and 5% $CO_2$ in a humidified incubator. Cell lines at confluency, moved the cells to the confocal live stage. Live stage maintained the incubator condition as same as previous cultured condition. The cellular media were replaced through the flow out/in tube by syringe. Images were obtained same methods with 'Confocal Imaging and Analysis'. For hypoxia condition, the live stage incubator $O_2$ concentration was regulated from 20% to 1%. Cells were in the 1% hypoxia condition at least 2 h for fully consumed the media and intracellular oxygen.

**Western Blotting**. For all cell lines used in this work, the cells were lysed under the appropriate conditions per experiment (such as FCCP treatment or 43 °C heat shock). Cells were lysed in PBS buffer containing 2% SDS (D0996, TCI) and 1× protease inhibitor cocktail (Invitrogen, 78438), for 5 min at 4 °C. Lysates were transferred to 1.5 ml tubes and cleared by ultrasonication (QSONICA, Q800R3) for 6 min in a cold-water bath.

Lysates were then centrifuged at 15,000 × *g* for 10 min at 4 °C, and the supernatants were collected. Lysate samples were boiled at 95 °C for 5 min in a 1x SDS-PAGE loading buffer. Subsequently, the proteins were loaded on 8% or 10% bis-tris gels for SDS–PAGE (Invitrogen) and subjected to immunoblotting with the indicated antibodies. The primary antibodies used for western blotting were anti-HSF1 (1:500 dilution, CST, Cat# 4356) and anti-Histone 3 (1:3000 dilution, Santa Cruz, sc-517576). Secondary antibodies, including anti-mouse (1:3000 dilution, CellNest, Cat# CNG004-0005) and rabbit-horseradish peroxidase (1:3000 diltution, CST, Cat# 7074S) were purchased from Invitrogen. For APEX2 samples, whole cell lysates (10 μl) were separated on a 10% SDS-PAGE gel. For blotting analysis, gels were transferred to nitrocellulose membranes (PALL, 0.2 μm), stained with Ponceau S (10 min in 0.1% (w/v) Ponceau S in 5% acetic acid/water), and then blocked in a solution of 2% (w/v) dialyzed BSA and 0.1% Tween-20 in Tris-buffered saline (TBST) at 4 °C overnight, or at room temperature for 1 h. The blots were immersed in streptavidin-horseradish peroxidase in blocking buffer (1:10,000 dilution, Thermo Scientific, Cat#21126) at room temperature for 30 min and then rinsed with blot blocking buffer three times for 5 min, before development using the Clarity reagent (Bio-Rad) and imaging on a LAS 4000 camera system. For assessing comparative enzyme expression levels, the membrane was stripped with a stripping buffer (100 mM 2-mercaptoethanol, 2% SDS, 62.5 mM Tris-HCl, pH 6.8). Blots were washed three times, 5 min per wash, in TBST buffer and incubated in a 5% blocking buffer for 1–2 h. They were then incubated in 10 mL of primary antibody solution, mouse-anti-V5 (1:5000 dilution, Invitrogen, Cat# R960-25),  for 1–2 h at room temperature. Blots were subsequently washed 3 × 5 min in TBST prior to incubation with 10 mL of secondary antibody, anti-mouse-HRP (1:3000 dilution, CellNest, Cat# CNG004-0005), diluted in blocking buffer, for 30–60 min at room temperature. The membranes were then washed again for three times, 5 min per wash in TBST buffer before developing using the Clarity reagent (Bio-Rad), as described above.

**APEX2 reaction with Endogenously generated ROS.** HEK293T cells were cultured in same condition as described in "Cell Culture". After 24 h of transfection, 250 μM DBP in DMEM was treated to the APEX2-expressed cells for 30 min, and then, 30 μM menadione was added for an appropriate time. The cells were washed with DPBS containing 10 mM Trolox, 5 mM sodium azide, and 20 mM sodium ascorbate three times, and RIPA lysis buffer (Elpis Biotech, Lexington, MA, USA, EBA-1149) was added after removing DPBS. The cells were lysed for 10 min at 4 °C, and the sample was loaded onto a 10% sodium dodecyl sulfate-polyacrylamide gel electrophoresis gel for separation at 220 V for 60 min. After separation, proteins on the gel were transferred to a nitrocellulose membrane at 30 V for 180 min. The protein loading level was checked by Ponceau staining; the stain was removed by washing with 1× TBST. Blocking was performed with 2% skim milk in TBST for 1 h. The blocking solution was replaced by incubating the membrane with the primary antibody (described above) in 2% skim milk for 1 h. After 3 washes (each for 5 min) with 1× TBST, the membrane was incubated with the secondary antibody in 2% skim milk in TBST for 1 h. After 3 washes with 1× TBST buffer, the signals were developed with an ECL kit (Bio-Rad, Hercules, CA, USA, 1705061), and images were acquired with a gel doc machine (cytiva, Amersham™ ImageQuant™ 800).

**Luciferase assay.** HEK293T cells were cultured in same condition as described in "Cell Culture". After 24 h of transfection (Luc+ and renilla plasmid mixture 1:1), cells in stressed condition for 1 h and 3 times DPBS wash. Prep the cells with DPBS and centrifuged 300 G, 1 min, 4 °C and remove the supernatant.

Reagent preparation

- 1X PLB: 5X Passive Lysis buffer (PLB) dilute with DW
  store at 4 °C (Max 1 month)
- LARII: resuspend the Luciferase Assay Substrate in Luciferase Assay Buffer II (LARII)
  store at -20 °C (Max 1 month) or -70 °C (Max 1 year)
- Stop & Glo reagent: add 2.1ml 50X S&G substrate to 105ml S&G buffer in the amber S&G reagent bottle
  vortex 10 seconds
  store -20 °C for 15 days
- for smaller amount of 1X S&G reagent: to the required amount of S&G buffer,
  add 50X S&G substrate to final 1X conc.

Dispense the recommended volume of 1X PLB into each culture vessel. Pipetting over 10 times and leave the tube 10 min at RT (cell lysate prepared). Dispense the 100ul LAR II in 96-well white plate (Corning, 354620). Dispense the 20ul PLB lysate in each well with multi pipet and pipetting 1 time. Measure firefly luciferase activity by SpectraMax i3x (Molecular Devices). Dispense the 100ul Stop & Glo reagent in each well with multi pipet and pipetting 1 time measure the Renilla luciferase activity.

**ChIP and ChIP-seq Library Construction**. ChIP experiments were performed as previously described (2021 NAR Cancer, Hwa-Ryeon Kim et al.) Briefly, a concentration of $3 \times 10^7$ single trypsinized HEK293T cells was used. The trypsinized cells were crosslinked with 1% formaldehyde, followed by quenching in 0.125 M glycine, and washed with PBS. The cell pellets were lysed with 1200 µl of cell lysis buffer [10 mM Tris-Cl pH 8.0, 10 mM NaCl, 0.2% NP-40] supplemented with protease inhibitor cocktail (cOmplete™ Protease Inhibitor Cocktail, Roche, Cat#11697498001) and 1 mM DTT. Chromatin was isolated by centrifugation at 7,400 rpm for 30 s. The pellet was gently resuspended in 1200 µl of nuclei lysis buffer [50 mM Tris-Cl pH 8.0, 10 mM EDTA, 1% SDS] containing protease inhibitor and 1 mM DTT. The chromatin lysate was sonicated (Diagenode, Bioruptor pico) for 10 cycles (30 s, and 30 s off), and the sonicated chromatin mixture was then incubated for 1 h with 60 µg of rabbit IgG, and 60 µl of Protein A magnetic beads (Invitrogen Cat#

10001) for pre-clearing. Immunoprecipitation was conducted with 6 ml of pre-cleared chromatin, 30 µl of the HSF1 antibody (CST, Cat# 4356), and 60 µl of Protein A magnetic beads overnight at a 4 °C rotator. The following day, the immune complexes were washed with IP Wash I Buffer, twice with high-salt buffer, once with IP Wash II buffer, and finally twice with TE (pH 8.0). The washed immune complexes were eluted by incubation with 200 µl of elution buffer [1% SDS and 0.1 M $NaHCO3$] for 30 min at a 45 °C thermomixer, shaking at 1,000 rpm. The eluate was de-crosslinked with RNase A (1 µg/µl) and 0.25 M NaCl and incubated overnight at a 65 °C water bath. The next day, the eluate samples were incubated for 2 h in a Proteinase K (NEB, Cat# P8107S) solution, and the immunoprecipitated DNA was purified with a QIAquick PCR purification kit (QIAGEN, Cat# 28106;) in 50 µl of EB (elution buffer).

The ChIP-seq libraries were constructed using 40 µl of purified ChIP DNA and NEXTflex$^{TM}$ ChIP-seq kit (PerkinElmer, Cat# NOVA-5143-02) according to the manufacturer's instructions. Briefly, ChIP DNA was end-repaired and size-selected (250–300 bp) using AMPure XP beads (Beckman, Cat#A63881). All subsequent procedures, from adenylation to PCR amplification, were performed according to the ChIP-seq library construction steps. The quality of the ChIP-seq libraries was determined by a Bioanalyzer using the High Sensitivity chip (Agilent), and the average size of the ChIP-seq libraries ranged from 250 to 350 bp. For multiplexing, equal molar quantities of libraries were combined by considering the sequencing depth per sample (20–40 million reads per library). The ChIP-seq libraries were sequenced using the Illumina NextSeq platform with single-end reads of 76 bases.

**RNA-seq Library Construction**. Total RNA was extracted using the QIAzol reagent (QIAGEN, Cat# 79306) according to the manufacturer's instructions. RNA-seq libraries were constructed using 5 µg of purified RNA and a NEXTflex$^{TM}$ Rapid Directional mRNA-seq kit (Cat# NOVA-5138-11; PerkinElmer). Briefly, purified RNA was poly-A selected and fragmented using a fragmentation enzyme. After first- and second-strand cDNA synthesis from a fragmented RNA template, adenylation for PCR amplification was performed according to the RNA-Seq library construction steps.

**RNA-seq.** The RNA-seq libraries were prepared with NEXTflex$^{TM}$ Rapid Directional mRNA-seq kit (catalog no. NOVA-5138-11, PerkinElmer). Total 10 µg of RNA was extracted using QIAzol (catalog no. 79306, QIAZEN) and isolated into mRNA through poly-(A) selection step. The mRNA was fragmentated and synthesized into cDNA. Adenylation and adapter ligation were performed, followed by PCR amplification to construct the RNA-seq libraries.

**Data analyses of RNA-seq.**

**(1) Identification of DEGs.** Raw reads from RNA-seq were aligned to the human reference genome assembly (mm9) using STAR mapping tool. The relative transcript abundances were measured in Reads Per Kilobase of transcript per Million mapped reads (RPKM) from Cufflinks. Among 200 genes which are up-regulated by TNF-α treatment, 128 genes were ranked by their mean log2 fold change between the 4 experimental groups of interest.

**(2) GSEA analysis.** GSEA was performed according to the instructions. The MSigDB v7.0 and reference database were used for transcriptional difference between each treatment.

**(3) Gene ontology (GO) analysis of genes.** A list of the defined 200 genes which were up-regulated by TNF-a treatment was used as an input for GO analysis with AmiGO tool.

### Statistics

Statistical analyses were performed using excel. The probability values (P-values) were obtained using a student's T test. The P-values lower than 0.05 ($P < 0.05$) were considered statistically significant, and the following symbols were used: * ($P < 0.05$), ** ($P < 0.01$), *** ($P < 0.001$), and **** ($P < 0.0001$).

**Plasmids Information**

| Name (expected size) | Features | Promotor/ Vector | Details |
|---|---|---|---|
| HSF1-linker-EGFP | *BsiWI*-HSF1-*HindIII*-13 a.a linker-*NotI*-EGFP-*STOP-BlpI-EcoRI* | UBC / pFUGW | 13 a.a linker: GAPGSAGSAAGSG (Patel, Avinash, et al. *Cell*, 2015) |
| Mito-V5-APEX2 | *KpnI*-Mito-*BamHI-NheI*-V5-APEX2-*Stop-NotI* | CMV/ pcDNA5 | Mito-: MLATRVFSLVGKRAISTSVCVRAH (matrix targeting sequence, Fornuskova et al., 2010) |
| SCO1-V5-APEX2 | *KpnI*-SCO1-V5-APEX2-*Stop-NotI* | CMV/ pcDNA5 | SCO1: IMS marker protein |
| TDRKH-V5-APEX2 | *HindIII-KpnI*-TDRKH-*BamHI-NheI*-V5-APEX2-*STOP-NotI-XhoI* | CMV/ pcDNA5 | TDRKH: OMM marker protein |
| GBP-V5-APEX2 | *AflII-HindIII-KpnI*-GBP-*BamHI*-V5-APEX2-*STOP-NotI-XhoI* | CMV/ pcDNA5 | GBP: Cytosol targeting protein |
| pMDLg/pRRE | | CMV / pMD | 3rd generation lentiviral packaging plasmid; Contains Gag and Pol; also requires pRSV-Rev (Addgene#12253) and envelope expressing plasmid (Addgene#12259) |
| pRSV-Rev | | pREP / pRSV-Rev | 3rd generation lentiviral packaging plasmid; Contains Rev; also requires pMDLg/pRRE (Addgene#12251) and envelope expressing plasmid (Addgene#12259) |
| HSPD1/HSPE1 promoter sequence short ver | *KpnI*-HSPD1 promoter sequence short ver-*SacI*-Luc(+)-SV40 late poly (A) signal-Enhencer | HSPDE1/pGL3 enhencer | TGCGCTCGGCGAGACAGGTCGTCGGCGGCGAGTGA GGGACAGAGTGCAGGGCGCACACCGCAATGAGCCC GTGTCCCCTCCCTCCGCCTCTACTCCCGCCCCGCGG CACCGCGTGTGCAGGCAGCTCCCACCCACTTCCCGT CAGCCCGGGCCCTGCAATCTGCACACCCTGCGCGC GAGCCCCGCCCCTCCCTACCCGCGCAGGGTGTGCTA GCGCGCTCAGCCCTCTCCGGCCGGCTTAGTCTAGTT CCCGGGCCTCGCTCGGTTCCAGAACTTTCCAGAAA ATGCCGCGCTCCCTACGGCTCAAGGGTCAAATCGCG TCATTTCCGGGAGGGGACGAAGGGGTAGTTCTTTCA CCTCGGCTGGGCGCCTAGAAAAGCCTAGAAACAGC TCCTTTTTTCTTCCGCCTCCGAGTCTTCGCGTCAGCG TCCTGCGCAGGGCCCTTGGGGCGAATCGCGGTGCG CGTCGGGGCGACCGCCCTCCCTCCCTGGGAGGGGC GAGGGGGCTAGCGGCGACCGCTGGGGCGAGCGCGC CTGCGCGCTGGGTGATTTTTTCACGTGTCGCCAGGG CCGGACTGCGAG (HSPD/E1 promoter sequence short ver) |
| NTS-pHluorin2 | *KpnI*-NTS-*BamHI*-pHluorin2-Stop-*NotI* | CMV/ pcDNA5 | NTS:EFMPEPAKSAPAPKKGSKKAVTKAQKKDGKKRK RSRKESYSVYVYKVLKQVHPDTGISSKAMGIMNSFV NDIFERIAGEASRLAHYNKRSTITSREIQTAVRLLLPGE LAKHAVSEGTKAVTKYTSAK (nuclear targeting sequence) and pHluorin2 is from addgene (Plasmid #73794) |

The nuclear export sequence, NES (LQLPPLERLTLD), was derived from residues 6–17 of the HIV-1 Rev protein. Protein processed size during translocation was obtained by programs: ApE (https://jorgensen.biology.utah.edu/wayned/ape/)

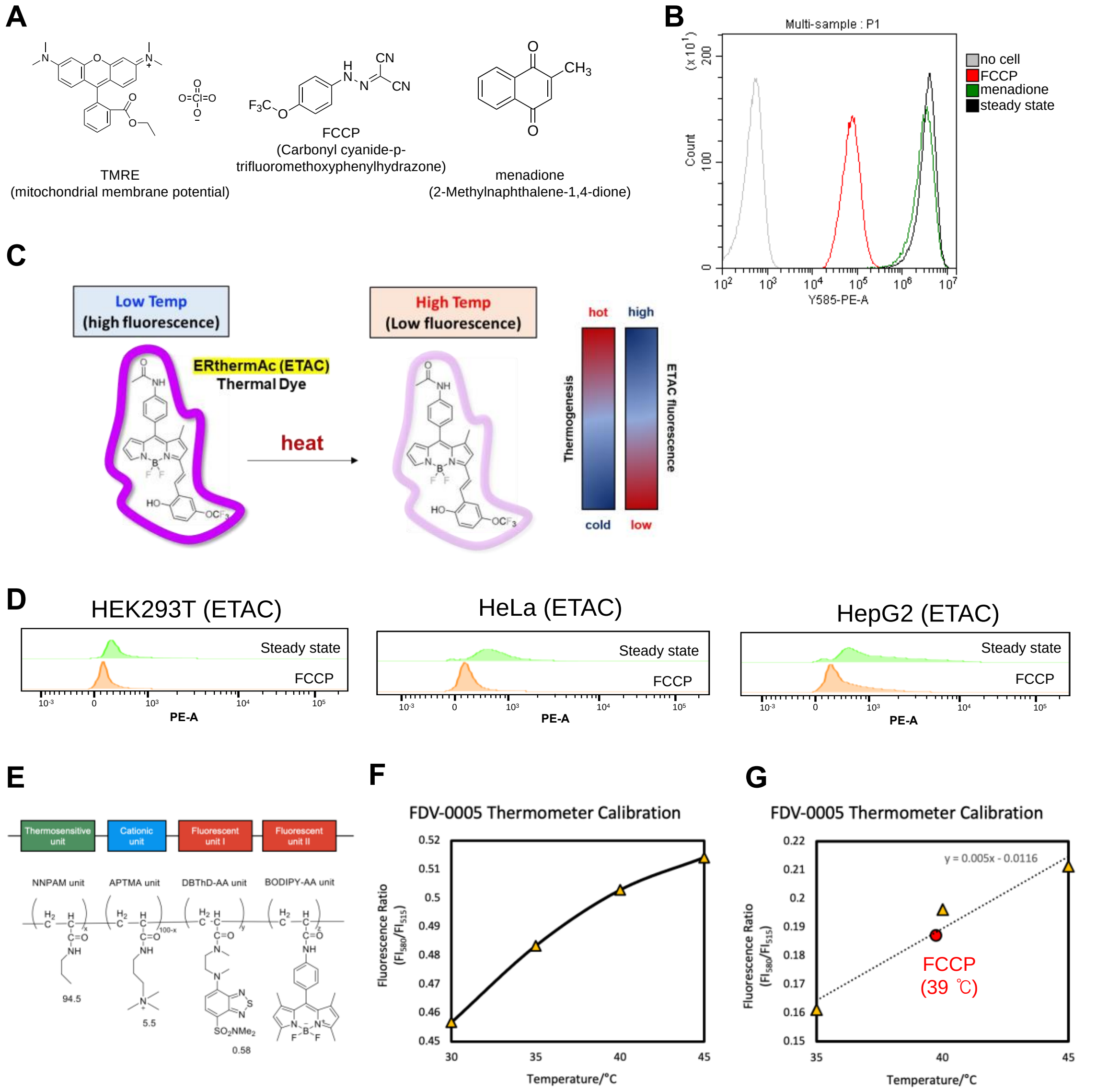


**Figure S1. Confirm the intracellular thermogenesis by two different thermometers.** (A) Chemical structures of TMRE, FCCP, and menadione. (B) Flow cytometry analyses and its histogram of HEK293T cell suspensions treated with either TMRE, FCCP, or menadione. (C) Schematic illustration of temperature measurement by ETAC. (D) ETAC fluorescence intensity after FCCP (100 µM, 1 h) treatment. Decreased ETAC fluorescence intensity under the indicates increased local temperature at the ER membrane. Flow cytometry analysis result of ETAC fluorescent signal under the two conditions (steady-state or 100 µM FCCP, 1 h), in HEK293T, HeLa and HepG2 cell lines. Cell count is shown on the y-axis, while fluorescent signal intensity (PE-A) is shown on the x-axis. (E) The polymer structure of the organic thermometer "FDV-0005". (F) Graph of the calibration curve and the FCCP treatment results for the organic thermometer. The thermometer was used at a dilution of 0.1% w/v in cell extract solution. Measurements were performed at a temperature range of 30 °C to 45 °C. (G) Intracellular temperature calibration by the FDV-0005 organic thermometer. The thermometer was used at a dilution of 0.01% w/v in 5% glucose solution. Measurements were performed at a temperature range of 35 °C to 45 °C. Cells were incubated at 37 °C and 5% $CO_2$ in a humidified incubator and treated with 100 µM FCCP.

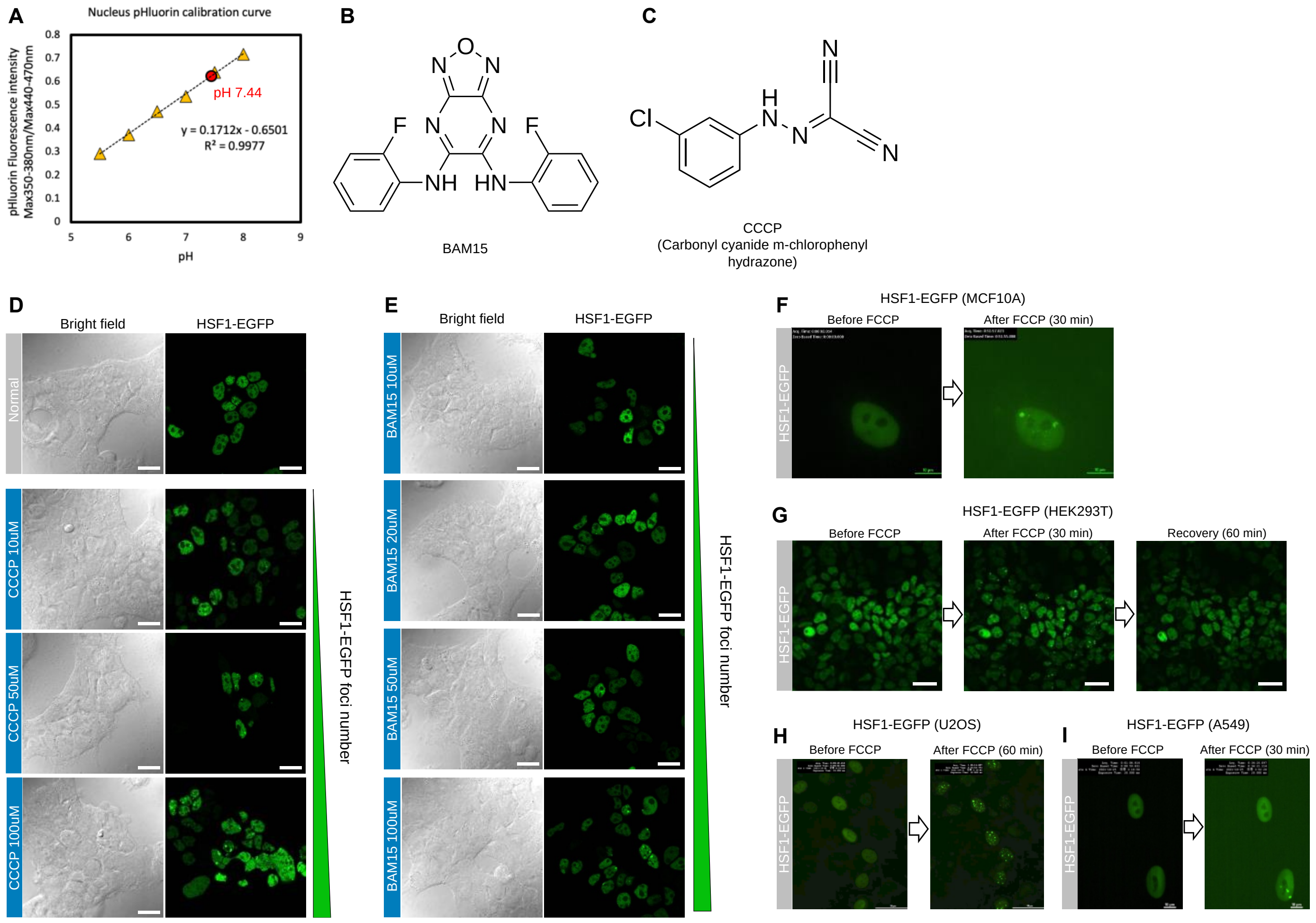


**Figure S2. Measurement of pH under FCCP condition and HSF1-foci formation in various HSF1-EGFP stably expressed cell lines.** (A) Measurement of nuclear pH in the FCCP condition. The fluorescent protein-based pH sensor pHluorin2 was used. The HEK293T cells were incubated at 37 °C and 5% $CO_2$ in a humidified incubator and 100 μM FCCP was used for this assay. (B-C) Chemical structures of BAM15 and CCCP. (D) Confocal images of HSF1-EGFP foci formation in HEK293T HSF1-EGFP-expressing cells following treatment with various CCCP concentrations (10, 50, and 100 μM; 1-h incubation). Scale bar 20 μm. (E) Confocal images of HSF1-EGFP foci formation in HEK293T HSF1-EGFP-expressing cells under treatment with various BAM15 concentrations (10, 20, 50, and 100 μM; 1-h incubation). Scale bar 20 μm. (F) Live-cell imaging of HSF1-EGFP foci formation following FCCP treatment (100 μM, 30 min) of HSF1-EGFP-expressing MCF10A cells. A real-time video recording is shown in movie 6. (G) Live-cell imaging of HSF1-EGFP foci formation following FCCP treatment (100 μM, 30 min) of HSF1-EGFP-expressing HEK293T cells. (H) Live-cell imaging of HSF1-EGFP foci formation following FCCP treatment (100 μM, 5 min) of HSF1-EGFP-expressing U2OS cells. A real-time video recording is shown in movie 5. (I) Live-cell imaging of HSF1-EGFP foci formation following FCCP treatment (100 μM, 30 min) of HSF1-EGFP-expressing A549 cells. All live cell experiments were conducted at 37 °C in a humidified 5% $CO_2$ incubator.

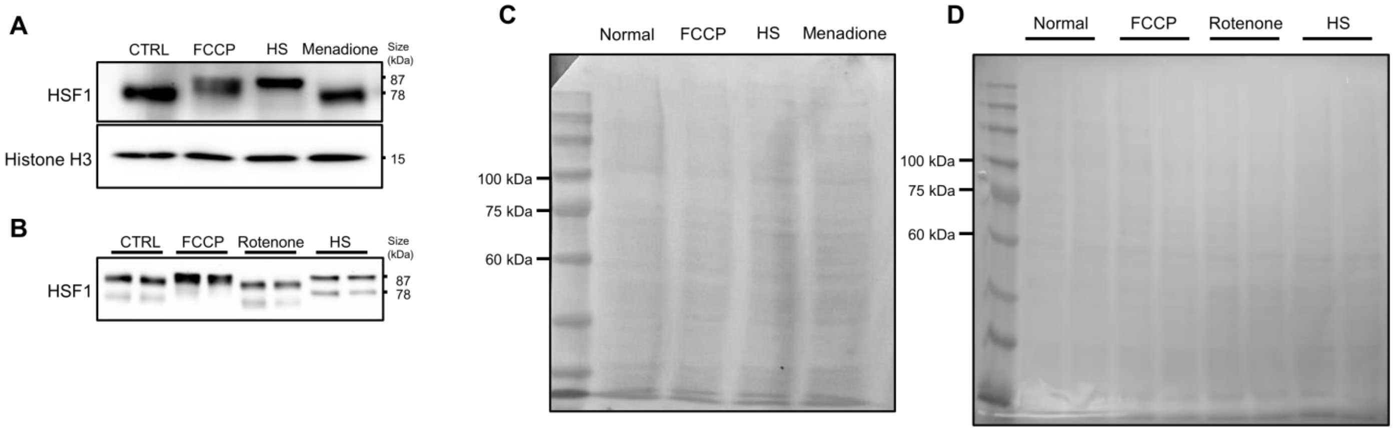


**Figure S3. Western blot analysis in MCF10A cells.** (A-B) Western blot analysis of (A) endogenous HSF1, or (B) recombinant HSF1-EGFP modification, following treatment with FCCP (100 μM, 1 h) or heat shock (43 °C, 1 h) in MCF10A cells. Histone H3 was used as reference. (C-D) Ponceau S image of western blot results (A) and (B) respectively.

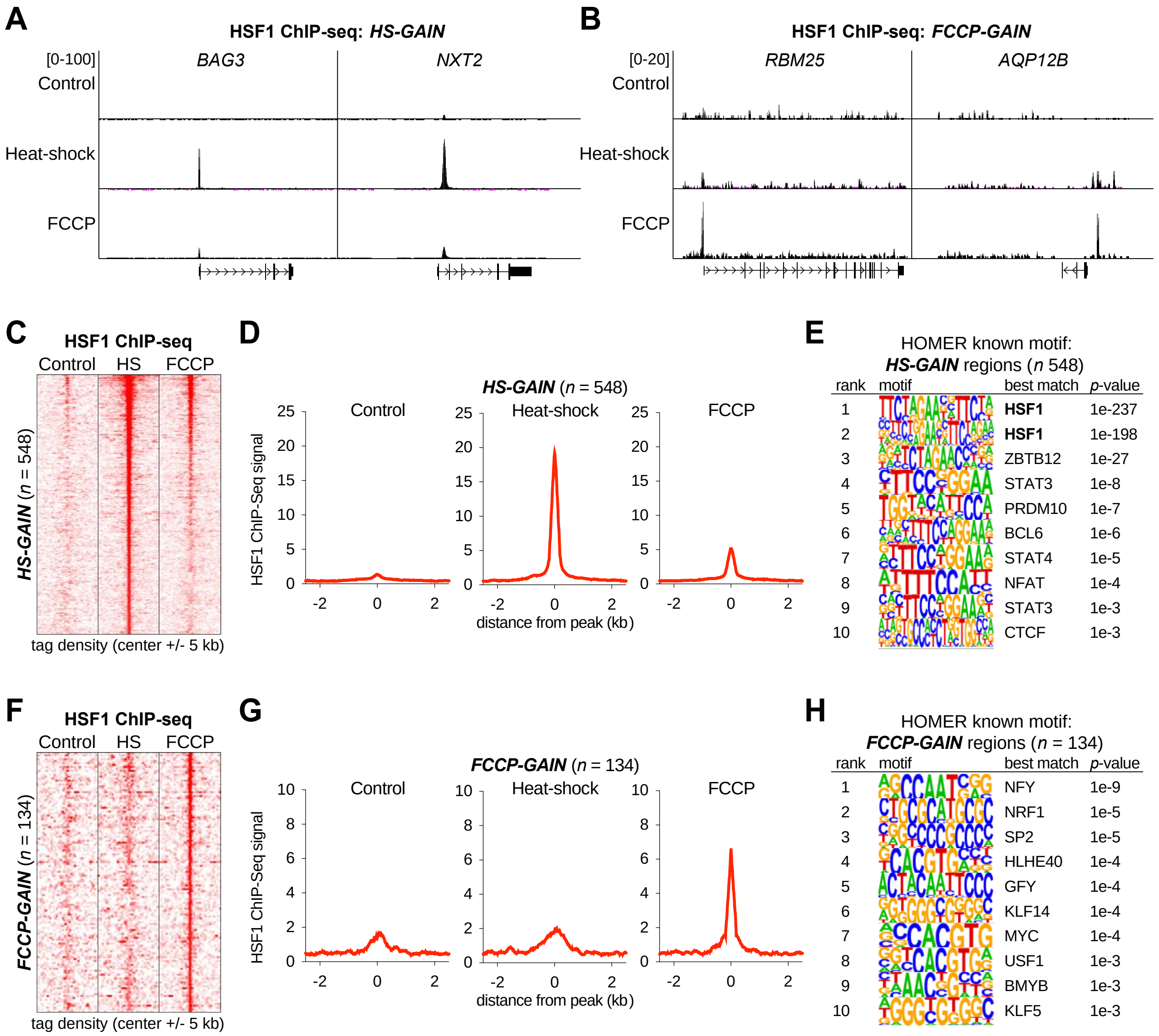


**Figure S4. Chip-seq and RNA-seq analysis of HSF1 under heat shock-specific and FCCP treatment-specific *GAIN*.** (A–B) Representative HSF1 ChIP-seq profiles at the (A) *BAG3* and *NXT2*, and (B) *RBM25* and *AQP12B* loci. (C) Density plots of HSF1 ChIP-seq, signal enrichment at a 5-kb region around the center of heat shock-specific bound regions. Each row represents a single region ($n$ = 548). (D) Analysis of the ChIP-seq signals (shown in C) overactive promoters and enhancers. The scale of all graphs was calculated using the distance of the peaks (kb). (E) Motif analysis (HOMER) based on HSF1 ChIP-seq results showing enriched motifs in *HS-GAIN* peaks. Only the top 10 statistically significant motifs are shown in the logo plots. (F) Density plots of HSF1 ChIP-seq and signal enrichment at a 5-kb region around the center of the FCCP treatment-specific bound regions. (G) Analysis of the ChIP-seq signals (shown in f) overactive promoters and enhancers. The scales of all graphs were calculated using the distance of the peaks (kb). (H) Motif analysis (HOMER) based on HSF1 ChIP-seq results showing enriched motifs in *FCCP-GAIN* peaks. Only the top 10 statistically significant motifs are shown in the logo plots.

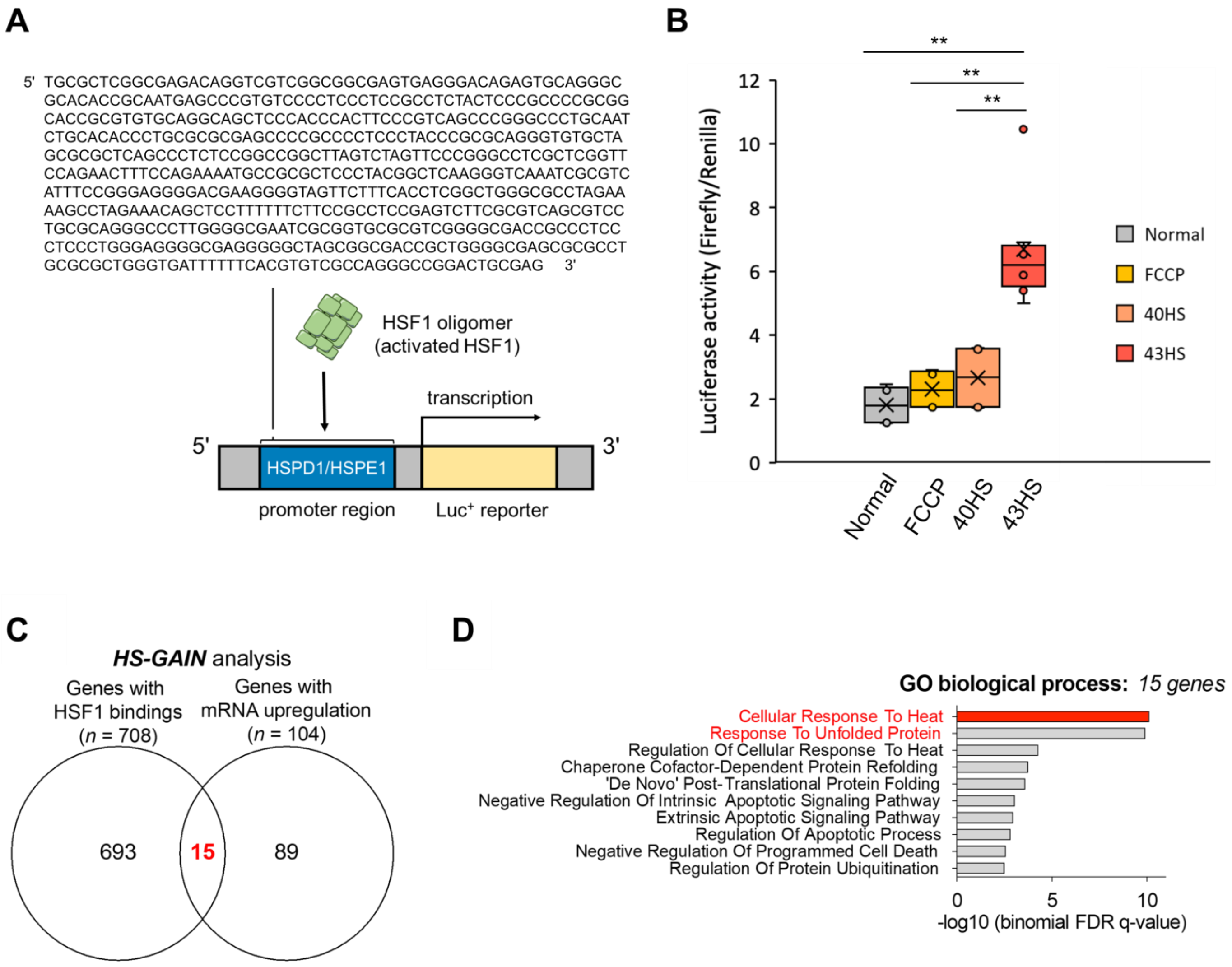


**Figure S5. Luciferase assay on *HSPD1*; HSF1 binding site and mRNA upregulated gene under Heat shock and FCCP treatment common-*GAIN*.** (A) HSPD1/HSPE1 promoter region sequence, and schematic representation of the HSPD1-luciferase assay. (B) HSPD1/HSPE1 promoter luciferase assay in HEK293T cells after FCCP treatment (100 µM, 1 h), heat shock at 40 °C for 1 h, or heat shock a 43 °C for 1 h). All live cell experiments were conducted at 37 °C, in a humidified 5% $CO_2$ incubator. Box plots shown in (B) indicate the quartiles, whiskers range from minimal to maximal values except outlier. The middle line of the box indicates the mid-point of the data (middle quartile). Each dot indicates individual data points plotted on the middle side of the box. Symbol 'x' means average value of data. Statistical analysis performed by student's T test and the following symbols were used:(**p<0.01). (C) Venn diagram showing the overlap ($n$ = 15) of the HSF1-bound genes ($n$ = 708) and mRNA upregulated genes ($n$ = 104) under heat shock-specific conditions. (D) Gene Ontology (GO) analysis of the biological processes associated with the identified 15 upregulated HSF1-bound genes (as shown in c).

**Supplemental Movie legends**

**Movie 1.** Crop of the Fig. 3B. U2OS cell. HSF1-EGFP stably expressed. 0 – 5 min: normal growth media, 15 – 40 min: FCCP media, 45 – 120 min: normal growth media

**Movie 2.** Wide view of Fig. 3B.

**Movie 3.** Crop of the Fig. S2F. MCF10A cell. HSF1-EGFP stably expressed. 0 – 30 min: normal growth media, 30 – 82 min: FCCP media

**Movie 4.** Wide view of Fig. S2F.

**Movie 5.** Movie of Fig. S2G. HSF1-EGFP for 30 min in normal growth media. HEK293T cell. HSF1-EGFP stably expressed.

**Movie 6.** Movie of Fig. S2G. HSF1-EGFP for 30 min in FCCP media. HEK293T cell. Continuing from Movie 5. HSF1-EGFP stably expressed.

**Movie 7.** Movie of Fig. S2G. HSF1-EGFP for 30 min in normal growth media. Continuing from Movie 6. HEK293T cell. Continuing from Movie 5. HSF1-EGFP stably expressed.

**Movie 8.** Movie of Fig. S2H. HSF1-EGFP foci formation in FCCP media. U2OS cell. HSF1-EGFP stably expressed. 0 – 30 min: normal growth media, 30 – 67 min: FCCP media

**Movie 9.** Wide view of Fig. S2H.

**Movie 10.** Movie of Fig. S2I. HSF1-EGFP foci formation in FCCP media. A549 cell. HSF1-EGFP stably expressed. 0 – 30 min: normal growth media, 30 – 35 min: FCCP media, 35 – 67 min: normal growth media

**Movie 11.** Movie of Fig. 4B. HSF1-EGFP foci segregated in FCCP media, 1% hypoxia condition. HEK293T cell. HSF1-EGFP stably expressed.
